# Comparison of the three-dimensional chromatin structures of adolescent and adult peripheral blood B cells: implications for the study of pediatric autoimmune diseases

**DOI:** 10.1101/2023.09.11.557171

**Authors:** Kaiyu Jiang, Yao Fu, Jennifer A. Kelly, Patrick M. Gaffney, Lucy C. Holmes, James N. Jarvis

## Abstract

**Background/Purpose:** Knowledge of the 3D genome is essential to elucidate genetic mechanisms driving autoimmune diseases. The 3D genome is distinct for each cell type, and it is uncertain whether cell lines faithfully recapitulate the 3D architecture of primary human cells or whether developmental aspects of the pediatric immune system require use of pediatric samples. We undertook a systematic analysis of B cells and B cell lines to compare 3D genomic features encompassing risk loci for juvenile idiopathic arthritis (JIA), systemic lupus (SLE), and type 1 diabetes (T1D).

**Methods:** We isolated B cells from healthy individuals, ages 9-17. HiChIP was performed using CTCF antibody, and CTCF peaks were identified. CTCF loops within the pediatric were compared to three datasets: 1) self-called CTCF consensus peaks called within the pediatric samples, 2) ENCODE’s publicly available GM12878 CTCF ChIP-seq peaks, and 3) ENCODE’s primary B cell CTCF ChIP-seq peaks from two adult females. Differential looping was assessed within the pediatric samples and each of the three peak datasets.

**Results:** The number of consensus peaks called in the pediatric samples was similar to that identified in ENCODE’s GM12878 and primary B cell datasets. We observed <1% of loops that demonstrated significantly differential looping between peaks called within the pediatric samples themselves and when called using ENCODE GM12878 peaks. Significant looping differences were even less when comparing loops of the pediatric called peaks to those of the ENCODE primary B cell peaks. When querying loops found in juvenile idiopathic arthritis, type 1 diabetes, or systemic lupus erythematosus risk haplotypes, we observed significant differences in only 2.2%, 1.0%, and 1.3% loops, respectively, when comparing peaks called within the pediatric samples and ENCODE GM12878 dataset. The differences were even less apparent when comparing loops called with the pediatric vs ENCODE adult primary B cell peak datasets.The 3D chromatin architecture in B cells is similar across pediatric, adult, and EBVtransformed cell lines. This conservation of 3D structure includes regions encompassing autoimmune risk haplotypes.

**Conclusion:** Thus, even for pediatric autoimmune diseases, publicly available adult B cell and cell line datasets may be sufficient for assessing effects exerted in the 3D genomic space.

## 1 Introduction

B cells are known to play an important role in the pathobiology of a wide range of autoimmune diseases that can have or exclusively have a pediatric onset (1–5). However, a serious limitation to our study of these cells in pediatric disease is the relatively small number of cells in the circulation (B cells represent about 20% of the circulating lymphocytes) and the limited volume of blood that can be acquired from children. Further complicating the problem are the practical and ethical considerations of acquiring blood from healthy children to isolate B cells that can be used as control or reference data.

After the completion of the mapping of the human genome, the National Institutes of Health embarked on an ambitious effort to characterize the non-coding, functional elements (i.e., those elements composing 98% of the genome) in medically-important organisms, cell lines, and primary human cells. These two projects, the <u>Enc</u>yclopedia of (Functional) <u>D</u>NA <u>E</u>lements (ENCODE) (6) and Roadmap Epigenomics (7) have provided investigators with a treasure trove of information that can be used to inform our understanding of data derived from patients with a broad range of diseases. Unfortunately, these genomic reference data sets are lacking data from children, although fetal and cord blood samples are represented in the collection. This data gap has forced investigators of pediatric diseases to generate their own genomic data sets on an ad hoc basis (8), often at considerable expense. This barrier limits the number of patients who can be studied in research aimed at understanding genetic/genomic mechanisms of disease, as the costs of acquiring samples from both patients and controls as well as the procedures themselves (e.g., sequencing costs) must be included in limited research budgets. The perceived need for acquiring reference samples from healthy children, rather than using ENCODE and Roadmap Epigenomics reference sets, represents on “hidden tax” on pediatric investigators that may not be placed on investigators of adult diseases, who can use these reference data without any concerns about their suitability.

Despite the concerns about using adult data or human cell lines for pediatric genomic research, the differences between adult and pediatric peripheral blood cells have not been examined in detail. Zhao et al demonstrate differences in the genomic locations of CpG methylation sites when they compared adult and pediatric CD4+ cells (9). Beyond this single study, however, little else is known. Here, we compare the three-dimensional chromatin structures of adult and pediatric peripheral blood B cells and with the commonly used B cell line, GM12878. We demonstrate that the three-dimensional loop structures within these different cell types differ at fewer than 1% of shared locations across the genome. Furthermore, we show that these small differences are no greater within the regions that harbor risk SNPs for juvenile idiopathic arthritis (JIA), systemic lupus erythematosus (SLE), and type 1 diabetes (T1D) than in the broader genome.

## 2 Materials and Methods

### 2.1. Study subjects

Healthy children were recruited under a University at Buffalo IRB-approved protocol **(#**MODCR00007255) from the University at Buffalo Medical Doctors Pediatric Clinic. Four subjects were included in this study, 3 boys (ages 9, 16, and 16) and 1 girl (age 17). Informed consent was obtained from the parents of the children providing samples, and assent was obtained from each of the participating children.

### 2.2. Isolation and Preparation of B cells

Whole blood (total 24 ml) was drawn into 8 mL citrated Cell Preparation Tubes (Becton Dickinson, Franklin Lakes, NJ, USA). Specimen processing was started within one hour from the time the specimens were drawn. PBMC were separated from granulocytes and red blood cells by density-gradient centrifugation. B cells were purified from PBMC by negative selection using the StemSep™ Human B Cell Enrichment Kit (STEMCELL Technologies Inc., Vancouver, Canada) following the manufacturers’ instructions.

### 2.3. CTCF HiChIP

B cells (~2x10^6^ each sample) were fixed in 1% formaldehyde at room temperature for 10 min. The fixation reaction was quenched by adding glycine (final concentration of 125 mM) and incubated for 5 minutes at room temperature with rotation. Cells were pelleted at 500xg at room temperature for 5min. Cell pellet were rinsed once using 1ml PBS. HiChIP assays were performed as described in Mumbach et al, 2016 with some modifications. Briefly, nuclei were isolated from fixed B cells and subjected to *in situ* digestion using 200U MboI (NEB, R0147M) for 4 hours at 37 ºC. The restriction fragment overhangs were filled and labeled with dCTP, dGTP, dTTP and biotin-dATP using Klenow DNA polymerase I (NEB M0212L), followed by *in situ* proximity ligation at room temperature overnight. The chromatin was fragmented by 2-minutes sonication with the Covaris E220 system, then immunoprecipitated using CTCF antibody (Cell Signaling, 3418S) at 4ºC overnight. DNA was purified by Zymo DNA Clean & Concentrator kit (Zymo, D4003). Streptavidin C1 beads were used to capture biotin-labeled DNA fragments. The sequencing libraries were generated on the streptavidin C1 beads using Illumina Tagment DNA Enzyme and Buffer kit (Illumina, 20034198) and run on the Illumina Novaseq S4 flowcell.

### 2.4. Statistical Analyses

#### 2.4.1. Initial QC and read processing

Adapters were removed from sequenced fastq files by fastp (options: detect_adapter_for_pe -l 50 -x –g) (10). Samples 3 (246M reads) and 6 (263M reads) were downsampled to 200M reads using seqtk for downstream processing (https://github.com/lh3/seqtk). Reads were aligned to the human genome using Bowtie2’s *H. sapiens* GRCh38 no alt analysis set with a mapQ > 10 and the following options: for global alignment: --very-sensitive -L 30 --score-min L,-0.6,-0.2 --end-to-end –reorder, and for local alignment: --very-sensitive -L 20 --score-min L,-0.6,-0.2 --end-to-end –reorder (11, 12). Reads were then processed using a MboI hg38 restriction fragment bed file and the HiC-Pro analysis pipeline (see **Supplementary Table 1 and Supplementary Figure 1** for QC stats) (13). CTCF HiChIP read tracks were RPCG normalized for viewing.

#### 2.4.2. Pediatric sample CTCF peak calling

HiChIP peak sets were called within each sample separately using MACS2 with defaults, a genome size = 2913022398, and FDR q = 0.01 (14). Peaks were evaluated for known ENCODE blacklisted regions, but no overlap was found. A consensus peak set was generated from individual peak profiles using DiffBind (15) for reproducible peaks observed in all four samples.

#### 2.4.3. CTCF loop calling

CTCF loops were called within the pediatric samples using HiChipper (16)using three CTCF peak datasets: 1) CTCF peak set called within the pediatric samples using MACS2 described in 2.4.2. above, 2) the ENCODE GM12878 CTCF ChIP-seq peak set (reference epigenome ENCSR000AKB, GRCh38 accession ENCFF017XLW processed 2020-10-01, and 3) the ENCODE primary B cell CTCF ChIP-seq peak set from two adult females (reference epigenome ENCSR682AXR, GRCh38 accession ENCFF919YTB processed 2020-10-20). Differential loop calling was determined using DiffLoop (17) with a Mango q < 0.01. Anchors within 500 bp were merged into single peaks. Loops were then restricted to those called within 2 or more samples of each comparison group and PETS ≥ 2 in both comparison groups, similar to previously reports (17, 18).

#### 2.4.4. Gene set enrichment analysis (GSEA)

Loop anchors were annotated to gene transcript start sites (TSS) within 1M bp using the GenomicRanges library in R. A GSEA was then performed for the 500 genes with distances to TSS < 45kb using the MsigDB hallmark gene sets (19, 20). An FDR < 0.05 was used to determine pathways enriched for genes.

#### 2.4.5. Evaluation of TAD structures and the genes within them

Contact maps for viewing in Juicebox were created for the pediatric samples using HiC-Pro’s hicpro2juicebox.sh utility (21). Topologically associated domains (TADS) were determined visually and analyzed for *ENRAP2, IRF1*, and *IL6* JIA risk haplotypes as previously described (22).

## 3 Results

### 3.1. CTCF loop structures within pediatric primary B cells are similar to those in ENCODE GM12878 and adult primary B cell datasets

A total of 393,236 CTCF peaks (range: 101,230-311,719) were called in the four pediatric samples with strong correlation (r^2^>0.7) and an average peak size of 257.08 bp (**Figure 1A, Supplementary Table 2)**. A set of 40,782 peaks (consensus peak set) were observed in all four samples **(Figure 1B, Supplementary Table 3)**. This number of consensus peaks called in the pediatric samples was similar in size to the number of CTCF peaks identified in the ENCODE GM12878 dataset (n = 41,017), while a bit smaller than the primary B cell dataset (n=53,854), and had average and median peak sizes of 463.60 bp and 397.0 bp (compared to GM12878: 309 bp and 356 bp; ENCODE primary B cell dataset: 459 bp and 480 bp).

**Figure 1.**
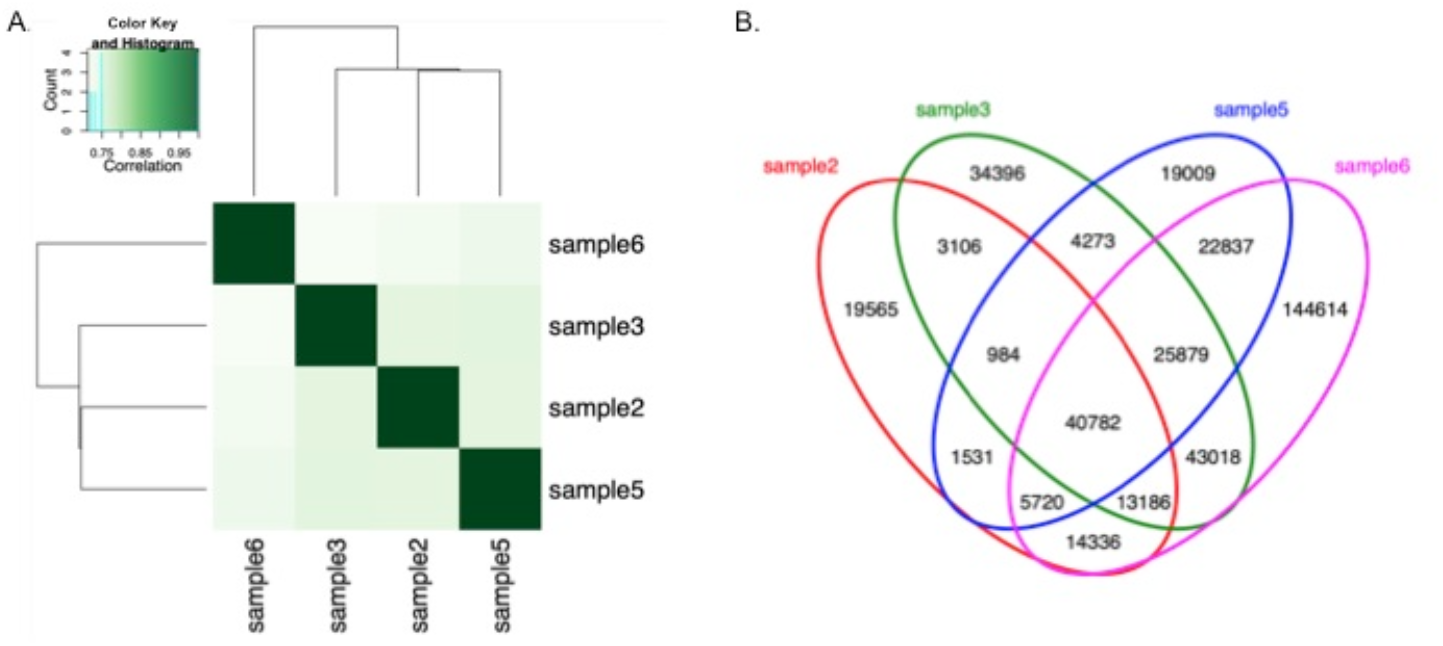
CTCF peaks in pediatric primary B cells. (A) Correlation heat map of CTCF peaks in four pediatric samples as called by MACS2. The four samples show strong reproducibility (r^2^ > 0.7) of peaks amongst them. (B) Venn diagram of shared CTCF peaks between the four pediatric samples as determined by DiffBind. A total of 40,782 peaks are observed in all four samples (consensus peak set).

When evaluating loops generated in the pediatric datasets using their own CTCF peaks and using the GM12878 CTCF peaks, a total of 61,860 shared CTCF loops (n = 181,316 prior to filtering) were called by DiffLoop for differential analysis (**Supplementary Table 4)**. Only 0.8% (n = 491) of loops were significantly different (FDR<0.05) between the two groups, of which 456 (93%) had significantly more PETs in the pediatric called peaks compared to the GM12878 peaks (**Figure 2A)**. The most significantly differential looping is on chromosome 22 between loop anchors at chr22:22679600-22688604 and chr22:22691237-22693712 with stronger loops observed in the samples when called with the pediatric peaks (**Supplementary Table 4; Supplementary Figure 2)**. This suggests that the overall 3D chromatin architecture in primary B cells collected from pediatric samples is similar to that of the publicly available GM12878 cell line dataset and that the cell line data should be sufficient for assessing effects exerted in the 3D genomic space of pediatric samples. The anchors that were involved in the 491 differential loops are near 661 genes (**Supplementary Table 4)**. Performing a GSEA showed that these genes are enriched (q < 0.05) in the following pathways: TNFA signaling via NFKB (q = 3.48E-5), allograft rejection (q = 1.07E-4), estrogen response early (q = 3.0E-4), myogenesis (q = 3.0E-4), and others (**Supplementary Table 5)**. The majority of genes (n = 489, 74%) were involved in only one differential loop (**Supplementary Figure 3a**). Several genes, however, were involved in multiple differential looping events, with the EH domain containing 1 gene (*EHD1*) on chromosome 11 being involved with the highest number of differential looping events: eight. However, this gene has not yet been associated with any trait or phenotype (https://www.ebi.ac.uk/gwas/search?query=EHD1). We provide a table of the genes involved in differential looping events in **Supplementary Table 6**.

**Figure 2.**
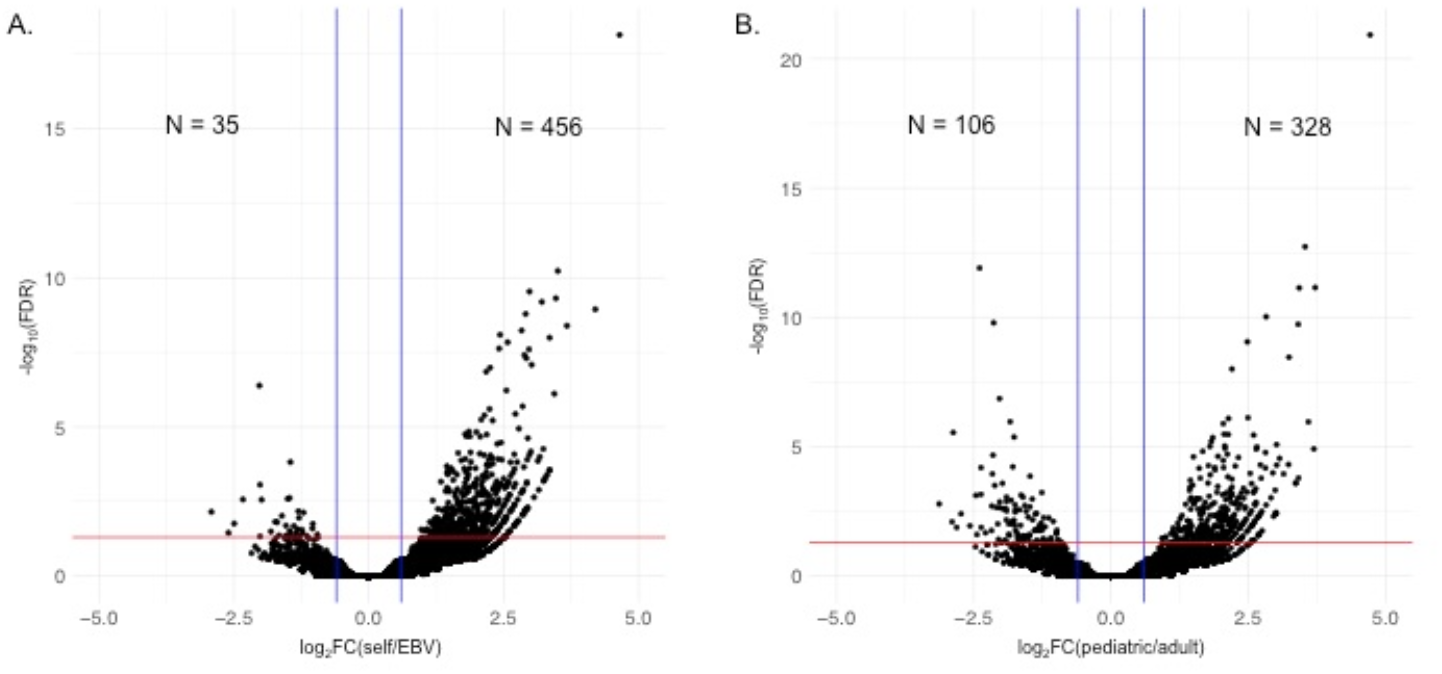
CTCF loop differential analysis volcano plots in pediatric samples. A. Differential analysis of loops using self-called pediatric CTCF peaks compared to ENCODE GM12878 CTCF peaks. B. Differential analysis of loops using self-called pediatric CTCF peaks compared to ENCODE adult primary B cell peaks. Vertical blue lines represent a FC ≥ 1.5 or ≤ -1.5 and red horizontal line represents an FDR q ≤ 0.05. Log_2_FC is plotted on the x-axis and –log_10_(FDR) is plotted on the y-axis. Numbers representing significantly higher PETs in self called peaks are to the right of the graph and those with higher PETs in ENCODE datasets to the left of the graph.

We next evaluated the ENCODE adult primary B cell CTCF peak dataset to see how similar it was to the peaks called in the pediatric primary B cell dataset. We observed fewer significant looping differences when comparing these two datasets than we observed when comparing primary pediatric and GM12878 cells. A total of 70,909 (n = 195,221 prior to filtering) CTCF loops were identified for differential analysis, with 434 (0.61%) of the loops showing significant differences (FDR<0.05) between the two groups (**Figure 2B, Supplementary Table 8)**. A total of 328 (75.6%) loops had significantly more PETs in the pediatric consensus peaks compared to the ENCODE adult peaks, while 106 loops had significantly more PETS in the ENCODE adult peaks. The 151 genes near anchors with more PETs in the ENCODE adult peak loops were enriched for the following pathways: myogenesis (q = 0.022), MYC targets version 2 (q = 0.022), coagulation (q = 0.022), apoptosis (q = 0.029), *IL2-STAT5* signaling (q = 0.042), and hypoxia (q = 0.042) (**Supplementary Tables 9A)**. The 459 genes near anchors with more PETs in the pediatric peak loops were enriched for the following pathways: hypoxia (q = 9.58E-5), *TNFA* signaling via *NFKB* (q = 2.92E-4), apical junction (q = 3.27E-3), *KRAS* signaling DN (q = 3.27E-3), *MTORC1* signaling (q = 3.27E-3), and others (**Supplementary Table 9B)**. Looking at the 591 total genes annotated near differential loops in this analysis, 455 (77%) were involved in only one differential loop (**Supplementary Figure 3B)**, and the greatest number of loops attributed to any gene was five.

### 3.2. Loop structures within and subsuming the JIA, T1D, and SLE risk haplotypes are not significantly different when using peaks called within pediatric primary B cells compared to publicly available datasets

We next sought to determine whether regions conferring risk for childhood-onset autoimmune diseases showed any notable differences in loop structures compared to the global analysis. We queried the risk regions for three autoimmune diseases in which B cells play an important role in pathobiology: JIA (5), T1D (1), and SLE (2). We first compared loops generated in the pediatric samples using the self-called MACS2 peaks to loops called in the pediatric samples using the ENCODE GM12878 B cell line peak dataset. A total of 313 CTCF-anchored loops (n = 1,119 prior to QC filtering) were shared in 29 regions conferring risk for JIA (**Supplementary Table 11)**. Only seven loops (2.2%) exhibited differential looping (FDR < 0.05) between the two groups, with all loops having more PETs in the pediatric called peaks. These loops were located in the following six JIA risk regions: *JAK1* (chr1:64661309-64851277), *AFF3-LONRF2* (chr2:100208154-100273161, and chr2:100208154-100273161), *WDFY4* (chr10:48581280-48686719), *ZFP36L1* (chr14:68790795-68851245), *TYK2* (chr19:10363179-10430870), and *IL2RB* (chr22:37139351-37210641). T1D risk haplotypes generated 1,350 shared loops (n = 4,423 prior to QC filtering) **(Supplementary Table 12)**. A total of 13 loops (1.0%) on 10 T1D haplotypes displayed differential looping, with all loops having more PETs in the pediatric called peaks. The most significantly differential loops were located on *IL2RB-C1QTNF6* haplotype (chr22:37139351-37210641 and chr22:37154012-37226586) **Supplementary Table 12)**. When looking at loops in regions conferring risk for SLE, a total of 2,398 loops (n = 7,532 prior to filtering) were shared (**Supplementary Table 13)**. Only 31 loops (1.3%) were significantly different between the two groups, with all loops having more PETs in the pediatric called peaks. These 31 loops involved 22 lupus haplotypes, with *ETV3-FCRL5, BLK*, and *IRF8* haplotypes containing more than one significantly differential loop **(Supplementary Table 13)**.

We then compared loops generated in the pediatric samples using the self-called MACS2 peaks to loops called in the pediatric samples using the ENCODE adult primary B cell peak dataset and observed even fewer differences between the two peak datasets. A total of 388 CTCF-anchored loops (n = 1,105 prior to QC filtering) were shared in 30 regions conferring risk for JIA (**Supplementary Table 14)**. Only five loops (1.3%) exhibited differential looping between the two groups. Two loops had significantly fewer PETs when called using the pediatric self-called peaks compared to the adult primary B cell peaks (*ATP8B2-IL6R:* chr1:154298690-154361733, and *ICAM3-TYK2* (chr19:10243816-10447681), while the remaining three differential loops had more PETs when called using the pediatric self-called peaks (all of which also had significantly more PETs in the self-called peaks when compared against the GM12878 peaks): *WDFY4* (chr10:48581280-48686719), *ZFP36L1* (chr14:68790795-68851245), and *TYK2* (chr19:10363179-10430870). T1D risk haplotypes generated 1,679 shared loops (n = 4,764 prior to QC filtering) **(Supplementary Table 15)**. A total of 10 loops (0.6%) on 9 T1D haplotypes displayed differential looping. Again, the two loops in the regions of *ATP8B2-IL6R* (chr1:154298690-154361733) and *ICAM3-TYK2* (chr19:10243816-10447681) had significantly fewer PETs in the self-called pediatric peaks compared to the adult B cell peaks while the remaining eight loops had significantly more PETs in the pediatric self-called peaks. When looking at loops in regions conferring risk for SLE, a total of 2,896 loops (n = 7,808 prior to filtering) were shared (**Supplementary Table 16)**. Only 27 loops (0.9%) on 15 lupus haplotypes were significantly different between the two groups; four loops had significantly fewer PETs in the pediatric self-called peaks compared to the adult primary B cell peaks (*PARP11:* chr12:3857402-3990865, *PLD2:* chr17:4705955-4930243, *ARID3A:* chr19:834045-1010097, and *TYK2:* chr19:10243816-10447681), and the remaining 23 loops had significantly more PETs in the pediatric self-called peaks. We provide two examples of JIA risk haplotypes that do not demonstrate differential looping (*IL2RA* and *JAZF1*) as well as an example with *ATP8B2*, which does produce significant differential looping between the pediatric self-called peak loops and the ENCODE data in Figure 3.

**Figure 3.**
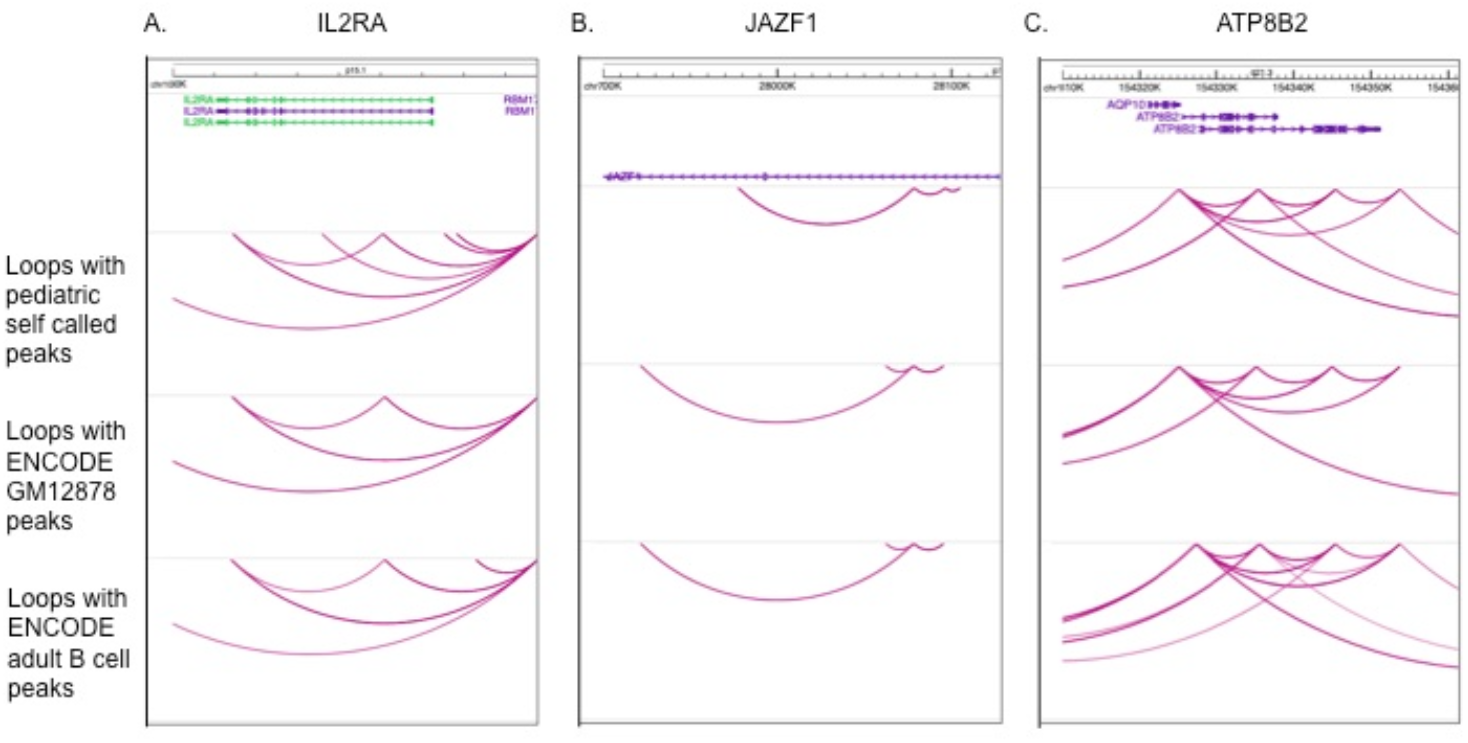
Illustrations of gene regions with similar and differential looping using CTCF peaks generated in pediatric B cell peaks and ENCODE datasets. A. *IL2RA* and B. *JAZF1* have similar loops when pediatric samples are generated with self-called CTCF peaks and ENCODE GM12878 and adult B cell peaks. C. *ATP8B2* has similar looping in pediatric B samples when self-called CTCF peaks are compared to ENCODE GM12878 peaks, but significant differential looping is observed when compared to ENCODE adult B cell peaks as determined by diffloop. Images were taken from the WashU epigenome browser. Loops were generated by HiChipper.

## 4 Discussion

As a group, the rheumatic diseases of childhood are among the most common chronic disease conditions in children (23, 24). B cells are known to play important roles in the pathobiology of these conditions (25–27), as well as other pediatric autoimmune diseases (3, 28). Furthermore, strong genetic risk associations have been identified for a broad spectrum of autoimmune diseases of pediatric onset. Because, for so many autoimmune diseases (29) including JIA (8, 30) and SLE (31) genetic risk is exerted most strongly by variants within the non-coding genome, elucidating genetic mechanisms will invariably require a broader understanding of the 3D genome of disease-relevant cells (8), with comparisons of cells from both healthy and affected individuals. The need for such comparisons was one of the important drivers of the ENCODE (6), Roadmap Epigenomics (7), and Blueprint Epigenomics (32) projects, which now provide invaluable reference data for the relevant epigenetic and functional features of the non-coding genomes of multiple tissues and cells. Whether these data sets can be used to interrogate genetic mechanisms in childhood onset diseases like JIA is unknown. Until now, the standard has been for investigators of these diseases to generate their own reference data from healthy children (8, 33, 34), something that is difficult and costly to do. The field of pediatric autoimmunity would benefit greatly if it could be ascertained that existing reference data sets are applicable, especially at autoimmune risk loci.

In this study, we used HiChIP to define the 3D architecture of pediatric B cells and compared findings with existing data sets derived from adult B cells and B cell lines. We find only small differences in the overall chromatin architecture when we compare consensus peaks between primary pediatric B cells and adult B cells and B cell lines. Furthermore, when we specifically examined the regions known to confer genetic risk for three autoimmune diseases (JIA, SLE, and T1D), we find no specific features that make the 3D chromatin architecture of these regions idiosyncratic. Therefore, for studies examining genetic mechanisms influencing B cells and driving these diseases, the existing reference data sets are, in most instances, likely to be suitable. Our findings therefore mirror those of Ray et al (35) who found that, for most studies aimed at elucidating genetic mechanisms in autoimmunity, the chromatin architecture of Jurkat T cells is sufficiently faithful to primary cells in most instances to allow their use. We would caution, as do Ray et al, that investigators carefully examine their locus of interest to be sure that the cell or cell line faithfully recapitulates primary cells in that region before undertaking mechanistic studies with CRISPR, CRISPRi, etc.

The discovery that much of the genetic risk for autoimmune diseases likely resides within the non-coding regulatory regions (8, 29) highlighted the importance of understanding the 3D structure of chromatin in disease-relevant cells. Regulatory structures such as enhancers may not regulate the most proximal gene in terms of linear genomic distance. Indeed, enhancers harboring disease-driving single nucleotide polymorphisms (SNPs) lying on autoimmune haplotypes may regulate genes that are not actually on the disease haplotype (36). However, enhancers and other regulatory elements typically regulate genes within the same chromatin loop or topologically associated domain (37). For example, in a genome-wide screening of 5,920 candidate enhancers, Gasperini and colleagues found that >70% of enhancers regulate genes within the same TAD (37). Thus, identifying target genes, i.e., the genes whose expression levels/function are influenced by the disease-driving SNPs on the autoimmune disease risk haplotypes, requires that one identify the relevant physical interactions between regulatory elements such as enhancers and the promoters of the candidate target genes. The arrival of newer methods to refine our knowledge of the 3D genome, such as MicroC, which can provide resolution at the nucleosome level (38) may greatly accelerate our ability to clarify genetic mechanisms by identifying target genes.

One limitation of this study is the fact that our pediatric sample consists of only one pre-adolescent, with the other B cell samples being taken from adolescents. It is possible that samples from younger, pre-pubertal children might display differences not seen in this mostly adolescent sample. The demographics of our sample reflect the challenges of obtaining peripheral blood cells from younger, healthy children. Given the small differences we observed in the 3 sample sets we studied here, however, there is reason to be confident that, for most queries into genetic effects exerted on the 3D genome, the data we have generated here will provide a suitable reference set.

In conclusion, we demonstrate that the 3D chromatin architecture of primary pediatric B cells differs little from that of adult B cells or B cell lines. Thus, for most studies at aim clarifying genetic mechanisms influencing gene expression in B cells, existing cell lines and reference data sets are likely to provide valid and relevant information. Furthermore, the data we have generated here will serve as a new and valuable reference data set for those regions where there may be identifiable and important differences between adult and pediatric B cells.

## 5 Conflict of Interest

The authors declare that the research was conducted in the absence of any commercial or financial relationships that could be construed as a potential conflict of interest.

## 6 Author Contributions

KJ: Writing – original draft, Investigation; YF: Writing – original draft, Investigation; JK: Writing – original draft, Formal analysis, Visualization; PG: Writing – original draft, Supervision; LH: Writing – review and editing; JJ: Writing – original draft, Conceptualization, Funding acquisition, Project administration, Supervision.

## 7 Funding

This work was supported by R03 AI166892, and R01AR078785 (to JNJ), and AR073606 and AI156724 (to PMG) from the National Institutes of Health. This work was also supported by the National Center for Advancing Translational Sciences of the National Institutes of Health under award number UL1TR001412 to the University at Buffalo. The content is solely the responsibility of the authors and does not necessarily represent the official views of the NIH.

## 8 Acknowledgments

Sequencing experiments were conducted by the Clinical Genomics Center at the OMRF (https://omrf.org/research-faculty/core-facilities/clinical-genomics-center/).

## Supplementary Material

**Supplementary Figure 1.**
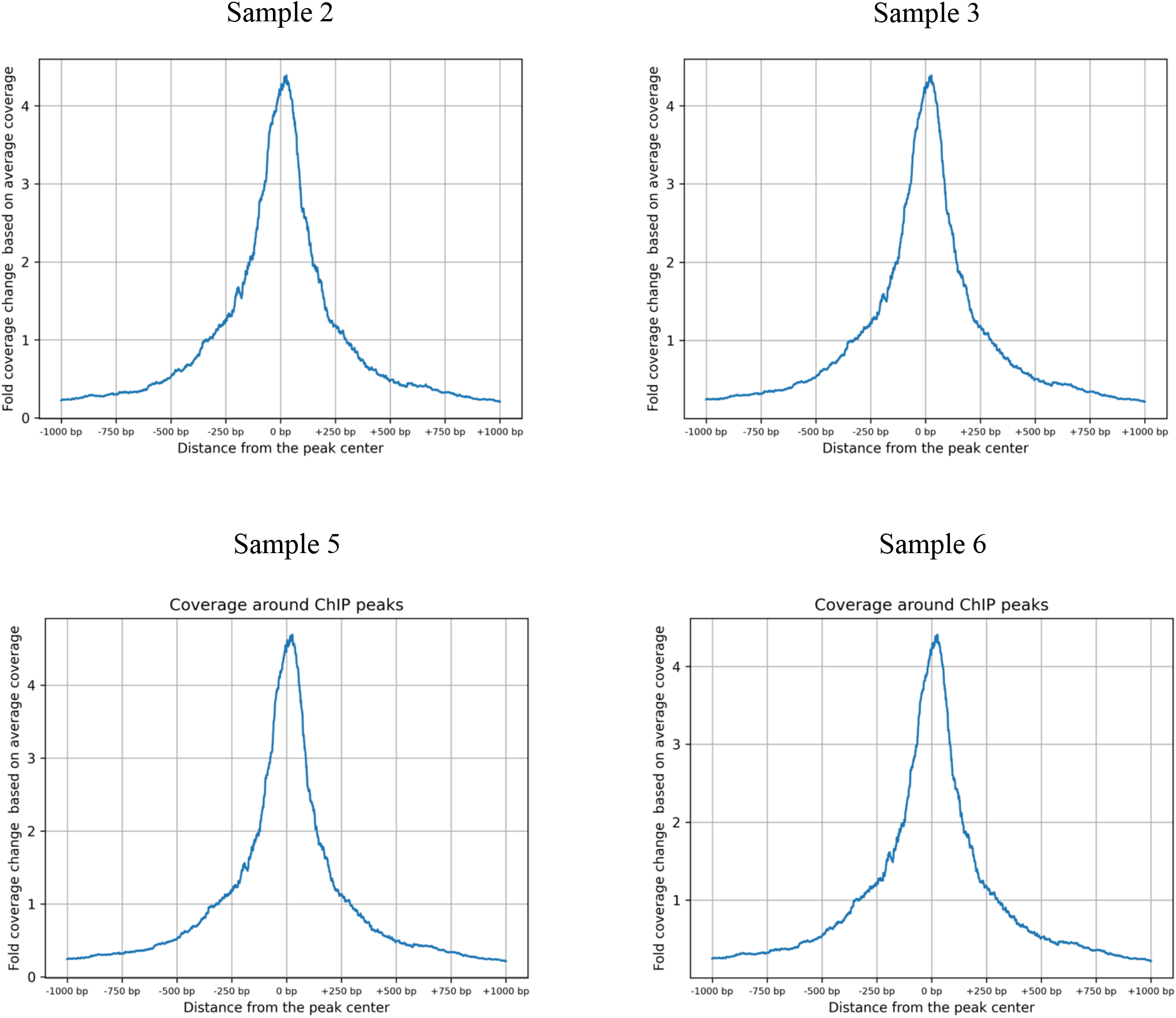
Global enrichment of reads around CTCF Hi-ChIP peaks in primary B cells collected from four pediatric samples. Distance from the peak center is plotted on the x-axis and fold coverage change based on average coverage is plotted on the y-axis. The fold coverage change based on average coverage is calculated by taking the mean HiChIP/base coverage at each base within 1kb of ChIp-seq peak center, then dividing the # read pairs at each base pair by the mean HiChIP/base coverage, and finally calculating the mean of the coverage fold change across all ChIP peak centers.

**Supplementary Figure 2.**
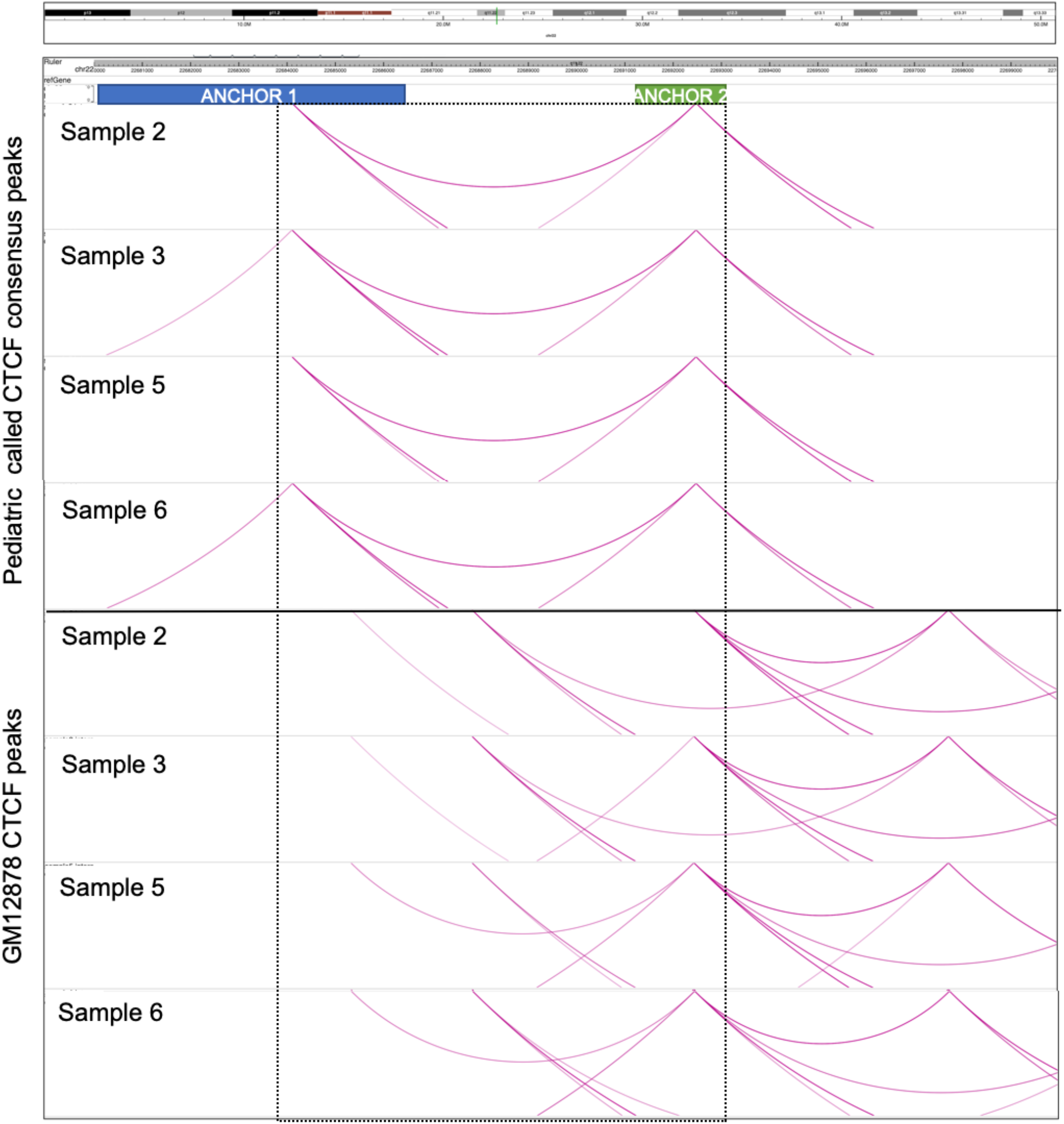
Most significant differential looping event between pediatric samples using self-called CTCF peaks or GM12878 CTCF peaks (chr22). Loops are shown for each pediatric sample with self-called peaks on top and GM12878 CTCF peaks on the bottom. The position of both anchors are shown and the loops attributed to the anchors are in the black square.

**Supplementary Figure 3.**
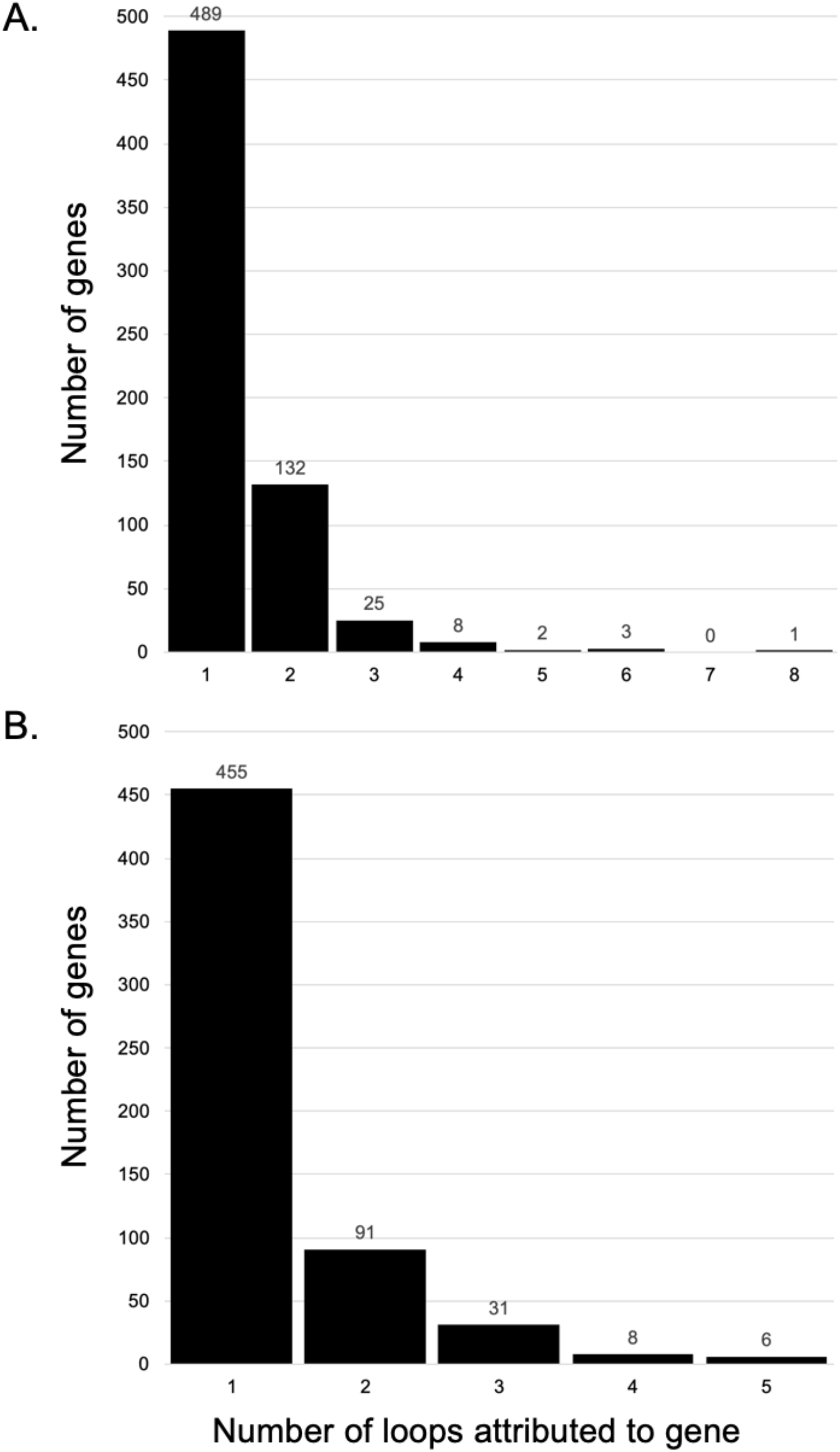
Number of significantly differential loops per gene. A. Pediatric primary B cell self-called CTCF consensus peaks and ENCODE GM12878 CTCF peaks. B. Pediatric primary B cell self-called CTCF consensus peaks and ENCODE primary B cell adult CTCF peaks.

## Notes

### Competing Interest Statement

The authors have declared no competing interest.

## References

1. Hinman RM, Cambier JC. Role of B lymphocytes in the pathogenesis of type 1 diabetes. Curr Diab Rep 2014;14:543.

2. Feng Y, Yang M, Wu H, Lu Q. The pathological role of B cells in systemic lupus erythematosus: From basic research to clinical. Autoimmunity 2020;53:56–64.

3. Arneth BM. Impact of B cells to the pathophysiology of multiple sclerosis. J Neuroinflammation 2019;16:128.

4. Piper CJM, Wilkinson MGL, Deakin CT, et al. CD19+CD24hiCD38hi B Cells Are Expanded in Juvenile Dermatomyositis and Exhibit a Pro-Inflammatory Phenotype After Activation Through Toll-Like Receptor 7 and Interferon-α. Front Immunol 2018;9:1372.

5. Wong L, Jiang K, Chen Y, Jarvis JN. Genetic insights into juvenile idiopathic arthritis derived from deep whole genome sequencing. Sci Rep 2017;7:2657.

6. ENCODE Project Consortium, Moore JE, Purcaro MJ, et al. Expanded encyclopaedias of DNA elements in the human and mouse genomes. Nature 2020;583:699–710.

7. Bernstein BE, Stamatoyannopoulos JA, Costello JF, et al. The NIH Roadmap Epigenomics Mapping Consortium. Nat Biotechnol 2010;28:1045–8.

8. Jiang K, Zhu L, Buck MJ, et al. Disease-Associated Single-Nucleotide Polymorphisms From Noncoding Regions in Juvenile Idiopathic Arthritis Are Located Within or Adjacent to Functional Genomic Elements of Human Neutrophils and CD4+ T Cells. Arthritis Rheumatol 2015;67:1966–77.

9. Zhao M, Qin J, Yin H, et al. Distinct epigenomes in CD4+ T cells of newborns, middle-ages and centenarians. Sci Rep 2016;6:38411.

10. Chen S, Zhou Y, Chen Y, Gu J. fastp: an ultra-fast all-in-one FASTQ preprocessor. Bioinformatics 2018;34:i884–i890.

11. Langmead B, Wilks C, Antonescu V, Charles R. Scaling read aligners to hundreds of threads on general-purpose processors. Bioinformatics 2019;35:421–432.

12. Langmead B, Salzberg SL. Fast gapped-read alignment with Bowtie 2. Nat Methods 2012;9:357–9.

13. Servant N, Varoquaux N, Lajoie BR, et al. HiC-Pro: an optimized and flexible pipeline for Hi-C data processing. Genome Biol 2015;16:259.

14. Zhang Y, Liu T, Meyer CA, et al. Model-based analysis of ChIP-Seq (MACS). Genome Biol 2008;9:R137.

15. Ross-Innes CS, Stark R, Teschendorff AE, et al. Differential oestrogen receptor binding is associated with clinical outcome in breast cancer. Nature 2012;481:389–93.

16. Lareau CA, Aryee MJ. hichipper: a preprocessing pipeline for calling DNA loops from HiChIP data. Nat Methods 2018;15:155–156.

17. Lareau CA, Aryee MJ. diffloop: a computational framework for identifying and analyzing differential DNA loops from sequencing data. Bioinformatics 2018;34:672–674.

18. Ji X, Dadon DB, Powell BE, et al. 3D Chromosome Regulatory Landscape of Human Pluripotent Cells. Cell Stem Cell 2016;18:262–75.

19. Subramanian A, Tamayo P, Mootha VK, et al. Gene set enrichment analysis: a knowledge-based approach for interpreting genome-wide expression profiles. Proc Natl Acad Sci U S A 2005;102:15545–50.

20. Mootha VK, Lindgren CM, Eriksson K-F, et al. PGC-1alpha-responsive genes involved in oxidative phosphorylation are coordinately downregulated in human diabetes. Nat Genet 2003;34:267–73.

21. Durand NC, Robinson JT, Shamim MS, et al. Juicebox Provides a Visualization System for Hi-C Contact Maps with Unlimited Zoom. Cell Syst 2016;3:99–101.

22. Jiang K, Kessler H, Park Y, Sudman M, Thompson SD, Jarvis JN. Broadening our understanding of the genetics of Juvenile Idiopathic Arthritis (JIA): Interrogation of three dimensional chromatin structures and genetic regulatory elements within JIA-associated risk loci. PLoS One 2020;15:e0235857.

23. Gortmaker SL, Sappenfield W. Chronic childhood disorders: prevalence and impact. Pediatr Clin North Am 1984;31:3–18.

24. Singsen BH. Rheumatic diseases of childhood. Rheum Dis Clin North Am 1990;16:581–99.

25. Wong L, Jiang K, Chen Y, Jarvis JN. Genetic insights into juvenile idiopathic arthritis derived from deep whole genome sequencing. Sci Rep 2017;7:2657.

26. She Z, Li C, Wu F, et al. The Role of B1 Cells in Systemic Lupus Erythematosus. Front Immunol 2022;13:814857.

27. Piper CJM, Wilkinson MGL, Deakin CT, et al. CD19+CD24hiCD38hi B Cells Are Expanded in Juvenile Dermatomyositis and Exhibit a Pro-Inflammatory Phenotype After Activation Through Toll-Like Receptor 7 and Interferon-α. Front Immunol 2018;9:1372.

28. Hinman RM, Smith MJ, Cambier JC. B cells and type 1 diabetes …in mice and men. Immunol Lett 2014;160:128–32.

29. Farh KK-H, Marson A, Zhu J, et al. Genetic and epigenetic fine mapping of causal autoimmune disease variants. Nature 2015;518:337–43.

30. Crinzi EA, Haley EK, Poppenberg KE, Jiang K, Tutino VM, Jarvis JN. Analysis of chromatin data supports a role for CD14+ monocytes/macrophages in mediating genetic risk for juvenile idiopathic arthritis. Front Immunol 2022;13:913555.

31. Hui-Yuen JS, Zhu L, Wong LP, et al. Chromatin landscapes and genetic risk in systemic lupus. Arthritis Res Ther 2016;18:281.

32. Adams D, Altucci L, Antonarakis SE, et al. BLUEPRINT to decode the epigenetic signature written in blood. Nat Biotechnol 2012;30:224–6.

33. Tarbell E, Jiang K, Hennon TR, et al. CD4+ T cells from children with active juvenile idiopathic arthritis show altered chromatin features associated with transcriptional abnormalities. Sci Rep 2021;11:4011.

34. Hui-Yuen J, Jiang K, Malkiel S, et al. B lymphocytes in treatment-naive paediatric patients with lupus are epigenetically distinct from healthy children. Lupus Sci Med 2023;10.

35. Ray JP, de Boer CG, Fulco CP, et al. Prioritizing disease and trait causal variants at the TNFAIP3 locus using functional and genomic features. Nat Commun 2020;11:1237.

36. Pelikan RC, Kelly JA, Fu Y, et al. Enhancer histone-QTLs are enriched on autoimmune risk haplotypes and influence gene expression within chromatin networks. Nat Commun 2018;9:2905.

37. Gasperini M, Hill AJ, McFaline-Figueroa JL, et al. A Genome-wide Framework for Mapping Gene Regulation via Cellular Genetic Screens. Cell 2019;176:377-390.e19.

38. Lee BH, Wu Z, Rhie SK. Characterizing chromatin interactions of regulatory elements and nucleosome positions, using Hi-C, Micro-C, and promoter capture Micro-C. Epigenetics Chromatin 2022;15:41.

